# Non-synaptic Mechanism of Ocular Dominance Plasticity

**DOI:** 10.1101/2025.09.02.673699

**Authors:** Maxwell K. Foote, William C. Huffman, Erin N. Santos, Philip R. Lee, Michal Jarnik, Wei Li, Juan S. Bonifacino, R. Douglas Fields

**Author notes:** **Corresponding author** R. Douglas Fields, PhD, National Institutes of Health, NICHD Bldg. 9, Room 1E126, MSC 0905, 9 Memorial Drive, Bethesda, MD 20892, and.

## Abstract

Classic experiments showing that monocular visual disruption alters synaptic connections to binocular neurons established the fundamental concept of synaptic plasticity. Synaptic inputs that are activated coincidently with postsynaptic action potential firing are strengthened, and inputs from cells firing before or after the postsynaptic action potential are weakened. An implicit assumption, however, is that the speed of impulse transmission is not altered by visual deprivation. If so, spike time arrival at binocular neurons would be affected, thereby inducing synaptic plasticity. This possibility is tested here in adult mice by monocular eyelid suture and monocular action potential inhibition in retinal axons. The results show that spike time arrival in visual cortex is altered by monocular visual disruption in association with morphological changes in myelin (nodes of Ranvier) on axons in optic nerve and optic tract. This non-synaptic mechanism of ocular dominance plasticity, mediated by myelin-forming cells, supplements and may drive synaptic plasticity.

## Introduction

A fundamental finding from the pioneering studies by Nobel Laureates Hubel and Wiesel (Wiesel and Hubel, 1963) is that monocular visual deprivation (MD) alters synaptic strength and connectivity in neurons that receive input from both eyes, strengthening input from the open eye and weakening synaptic input from the deprived eye. Binocular deprivation fails to alter synaptic strength, because there is no activity-dependent competition in spike time arrival favoring either eye (Wiesel and Hubel, 1965). This form of synaptic plasticity is driven by differences in synchrony of spike time arrival from the two eyes onto neurons that receive binocular input in the visual cortex.

This process applies more generally in other neural networks where multiple inputs converge onto a common postsynaptic neuron. This basic synaptic learning rule (Hebb, 1949) is termed spike-time dependent plasticity (STDP) (Bi and Poo, 1998; Markram et al., 1997). Synaptic inputs to neurons are weakened if they do not arrive simultaneously with action potential firing of the post synaptic neuron.

What has been overlooked is that if the speed of impulse transmission is altered from one eye by visual deprivation, then by changing the synchrony of spike time arrival, axonal plasticity would influence synaptic plasticity and ocular dominance shifts (Munyeshyaka and Fields, 2022).

Neural impulse transmission speed is determined in part by the morphology of electrogenic nodes of Ranvier (NOR) along myelinated axons (Waxman, 1980). Changes in NOR morphology (length of the nodal gap) and myelin thickness in optic nerve have been shown to alter neural impulse transmission speed, visually evoked spike time arrival in the visual cortex, and visual acuity (Dutta et al, 2018). This finding, in which nodal gap length was reversibly increased by inhibiting perinodal astrocyte exocytosis in transgenic mice, together with recent research on NOR plasticity in motor (Bacmeister et al., 2022) and spatial (Cullen et al., 2021) learning, and chronic stress (Miyata et al., 2016), requires tests to determine whether visual disruption changes conduction velocity and NOR structure in optic nerve and optic tract.

The present studies address visual system plasticity in adult mice for several reasons. Less is known about visual system plasticity in adults than during early postnatal development, even though visual impairment in adults is a substantial medical problem. Moreover, plasticity mechanisms in adulthood can be applied more broadly to other neural circuits in mature animals. Many developmental processes in the early postnatal period are altered by use-dependent effects, including cell proliferation, differentiation, survival (Priya et al., 2018), cell migration, neurite outgrowth (Okujeni and Egert, 2019), synaptogenesis (Yee et al., 2024), synapse elimination (Nelson et al., 1989), phagocytosis of synapses by microglia (Schafer et al., 2012), and more, which operate through different cellular mechanisms. For review see (Spitzer, 2006). Similarly, visual deprivation in early life has multifaceted effects on the development of oligodendrocytes which form myelin in the optic nerve and optic tract (Etxeberria et al., 2016) and in the developing cerebral cortex (Korrell et al., 2019). These diverse developmental effects interact in complex ways to influence myelination, resulting in contradictory experimental outcomes and ambiguity. For review see (Fields, 2013).

Furthermore, structural remodeling of compact mature myelin is a comparatively slow process, requiring a week or more (Dutta et al., 2018). In contrast, several mechanisms of synaptic plasticity proceed with much faster kinetics through changes in glutamate receptor expression (Philpot et al., 2001; Gu et al, 1989), inhibitory neuronal function (Hensch et al., 1989; Keck et al., 2011), and other physiological mechanisms that do not involve structural changes. Finally, the effects of prolonged visual deprivation have greater clinical relevance to ocular injury and disease than does brief monocular deprivation.

### Activity-Dependent Competition vs. Use/Disuse Effect

The effects of MD on NOR morphology during early postnatal development or in adults have not been investigated previously. It is important to distinguish between potential NOR plasticity being driven by a use/disuse effect or by a competitive process based on differences in the synchrony of action potential firing among axons.

Our previous research shows that 30 days of binocular visual deprivation (BD) in adult mice does not alter nodal gap length in optic nerve significantly (Santos et al., 2022) and does not affect recovery of nodal gap length after it is lengthened through inhibition of perinodal astrocyte exocytosis (Santos et al., 2024). These findings parallel synaptic plasticity in the visual system, which is driven by monocular, but not by binocular deprivation. The lack of NOR plasticity following BD indicates a lack of an observable activity-dependent trophic effect on nodal gap length.

Activity-dependent competition from the two eyes alters synaptic strength and connectivity in neurons receiving binocular input in visual cortex, but each optic nerve carries input from only one eye. Therefore, there is no activity-dependent difference in action potential firing among axons within optic nerve to drive a competitive process of NOR plasticity via MD. The optic tracts, however, (located beyond the optic chiasm where the nerves partially decussate) contain axons from both eyes, giving oligodendrocytes access to neuronal traffic from both eyes. Optic tract axons from the deprived eye are spontaneously active, but they lack the coordinated firing induced by vision. In contrast, axons in the same tract from the open eye convey correlated firing driven by visual experience. This anatomy presents a situation to test the hypothesis that an activity-dependent competition among axons from the open and occluded eye could alter NOR morphology in an axon-specific manner.

### Oligodendrocyte Mediated Plasticity

In contrast to synaptic plasticity, where coincidence of activity from the two eyes is detected by postsynaptic neurons, in NOR plasticity, spike synchrony in the optic tract must be detected by oligodendrocytes, which are situated along the axon, far from synaptic terminals.

Recently a theory of oligodendrocyte-mediated plasticity (OMP) has been proposed to address this problem (Pajevic et al., 2023). A single oligodendrocyte forms myelin through up to 50 slender cell processes extending from the soma. Each process wraps a segment of myelin independently around a different axon. According to OMP theory, an oligodendrocyte acts as a coincidence detector of action potential activity among individual axons in a manner analogous to coincidence detection of synaptic inputs in postsynaptic neurons. The theory proposes that an oligodendrocyte alters myelin on each of the axons it myelinates to change conduction velocity to minimize delays of spike time arrival in each axon relative to the population of axons within its domain.

Mathematical modeling shows that the OMP mechanism greatly increases synchronization of action potential arrival at the target (Pajevic et al., 2023). When spikes in axons fire in a coherent manner synchrony of spike time arrival at the target is greatly improved, but there is no change when spike firing among axons is not coherent. In bundles of axons with mixed firing patterns, axons are sorted by their correlated firing patterns. This would predict an axon-specific change in NOR morphology in optic tract following MD that is specific to axons from the open and sutured eye in the same tract.

To determine whether visual deprivation alters NOR gap length in this manner in mature mice, monocular eyelid suture was performed on mice of both sexes at postnatal day 40 (P40) and NOR lengths on retinal ganglion cell (RGC) axons were measured in optic tracts by confocal microscopy at P70. The latency to peak visually evoked potential in visual cortex was measured to determine whether MD alters spike time arrival in the visual cortex.

Although there is no activity-dependent competition in optic nerve, because it lacks binocular input, if a difference in action potential firing were imposed in optic nerve axons by experimental intervention, then by hypothesis, NOR plasticity would be induced specifically on axons with normal or inhibited firing. This prediction was tested by inhibiting action potential firing in a subset of optic nerve axons by transfecting the inwardly rectifying potassium channel (Kir2.1) selectively into retinal ganglion axons of one eye to hyperpolarize those axons. This was accomplished by intraocular injection of AAV2-EF1a-FLEX-Kir2.1-T2A-tdTomato (Xue et al., 2014) and verified by electrophysiological measurements.

The results show that MD alters the latency of action potential arrival in visual cortex in accordance with changes in NOR gap length, and that the effects are axon-specific, as predicted by OMP theory. This previously unrecognized plasticity of NOR morphology and conduction velocity in the visual system must be taken into consideration in studies of MD and in the clinical setting. More generally, the findings support the existence of a form of extra-synaptic plasticity regulating spike time arrival that can occur in parallel with, or even drive, changes in synaptic plasticity in neural circuits.

## Materials and Methods

## Materials

### Mouse Strains

Wild-type mice of both sexes were acquired from Charles River (C57BL/6). To express the potassium channel Kir2.1 specifically in retinal ganglion cells, which express Vglut2+ transporter (Want et al., 2020), mice with a Vglut2-ires-cre knock-in were acquired from Jackson Laboratory (Vong et al., 2011; Pauli et al., 2022 ). All mice were housed in a 12-hr light/dark cycle with access to food and water ad libitum, following protocols approved by the NICHD/NIH Animal Care and Use Committee.

### Plasmid Construct and Virus Production

Plasmid construct EF1α-FLEX-Kir2.1-T2A-tdTomato was gifted (Zhang et al., 2019), which was previously created using pAAV-MCS and #60661 (Addgene) backbones (Xue et al., 2014). Plasmid DNA for packaging was obtained from Boston Children’s Hospital viral core facility. A viral construct was prepared by the NINDS Viral Production Core Facility, using an AAV serotype 2 vector encoding AAV2-EF1α-FLEX-Kir2.1-T2A-tdTomato, with a final titer of 7 µg.

### Antibodies

Primary antibodies used were: Rabbit anti-Red fluorescent protein (1:200, Abcam, 124754); Guinea Pig anti-RBPMS (1:100, Phosphosolutions, 1832); Rabbit anti-enhanced green fluorescent protein (1:1000, Abcam, 290); Mouse anti-Cre recombinase (1:500, Millipore, 3120); Mouse anti-CASPR/Neurexin IV (K65/35) (1:50, UC Davis/NIH NeuroMab Facility, 73-001); Rabbit anti-Sodium Channel 1.6 (1:100, ASC009); Secondary antibodies used from Thermofisher Scientific, raised in mouse, rabbit or rat dependent on host species of primary antibody, highly cross absorbed and conjugated to fluorophores of Alexa Flour 488, 568, or 633 and used at 1:1000 dilution. For retinal nuclear staining, we used DAPI and Hoechst Nucleic Acid Stains (1:1000, Thermofisher, 62248).

## Methods

### Monocular Deprivation

Mice were prepared the day before eyelid suture with carprofen for pain relief and continued for 2 days after the procedure. The adult mice at P40 were anestheticized with Ketamine (0.1 mg/g: NDC 11695-0703-1) and placed under a dissecting microscope. The eyes are treated with Proparacaine Hydrochloride Ophthalmic Solution (USP 0.5%: NDC 61314-016-01) for discomfort relief and Ciprofloxacin Ophthalmic Solution (USP 0.3%.: NDC 61314-656-05) for antibacterial treatment. One eyelid was sutured together using 6-0 nylon P-3 reverse cutting sutures (AD surgical) under sterile conditions. The other eye was left open. Triple Antibiotic Ophthalmic Ointment (TAOO) was used liberally on the sutures after surgery to prevent the eyes from drying out and to prevent infection. All ophthalmic solutions and chemical restraints used on animals were acquired through the NIH VDR. The mice were checked regularly to ensure that the eyelid suture was intact, and if not, the mice were excluded from further experimentation.

### Monocular Inhibition

Mice at postnatal day 40 were transfected under Ketamine (0.1 mg/g: NDC 11695-0703-1) and Xylazine (10 mg/kg NDC 59399-110-20) anesthesia and placed under a dissecting microscope. One eye of mice was injected with AAV2-EF1a-FLEX-Kir2.1-T2a-tdTomato at postnatal day 40. To accomplish this the eyes were treated with Proparacaine Hydrochloride Ophthalmic Solution (USP 0.5%: NDC 61314-016-01) to manage pain. Cyclopentolate Hydrochloride Ophthalmic Solution (USP 1%: NDC 61314-396-03) and Phenylephrine Hydrochloride Ophthalmic Solution (USP 10%: NDC 42702-103-05) were administered to dilate the pupil to aid in visualization of the injection behind the lens. Working under a dissecting microscope, one eye was proptosed, and a small opening was created on the side of the eye just above the retina using a 33-gauge beveled needle. The injection was done using a 38-gauge blunt needle inserted into the opening and guided behind the lens using the microscope to view the needle as seen through the pupil. AAV (2µL at 1x10^1^ vg/mL of EF1α-FLEX-Kir2.1-T2A-tdTomato or CTB 488 or CTB 555 to identify nodes of Ranvier by staining paranodes (2µL of 1 mg/mL, Fisher Scientific) were slowly injected at a rate of 200 nL/s using a microinjection syringe pump (World Precision Instruments catalog #Micro2T). After injection, the needle position was held in place for 30 seconds before being slowly removed. The area was treated liberally with TAOO (NDC 16571-754-53) to prevent the eyes from drying out, and to prevent infection.

Patterned electroretinogram responses (pERG) were measured 30 days post injection (dpi) and again 60 dpi to determine that the Kir2.1 gene was expressed and was inhibiting action potential firing until the end of the experiment on postnatal day100 as described below.

### Animal Perfusion and Tissue Preparation

Transcardial perfusion fixation, with 4% paraformaldehyde (PFA) in phosphate-buffered saline (PBS) buffer was used for confocal microscopy and 4% paraformaldehyde + 2.5% glutaraldehyde in cacodylate buffer was used for electron microscopy. After fixation, optic nerves and tracts for examination by confocal microscopy were dissected from the brain and placed into 4% PFA (Electron Microscopy Sciences) in PBS over night at 4°C and then transferred to 30% sucrose in PBS for 3 days. Optic nerve and optic tract issue were embedded in optimal cutting temperature (OCT) embedding medium (Fisher Scientific) sectioned on a cryotome into 12 µm-thick slices and mounted on glass slides. Slides were stored at -20°C and rehydrated with PBS for immunocytochemistry.

### Retinal Removal and Fixation

Immersion fixation with 4% PFA was used for analysis of retinas. Eyes were removed and transferred to a dish with a silicone gel bottom, and fine dissection scissors were used to cut a circular hole around the cornea, taking care to avoid cutting the retina. The cornea and lens were gently removed, leaving the retinal cup. Four to five symmetrical, radial cuts were made through the retina using microdissection scissors. The cuts were made about 2/3 of the way to the optic disc to allow the retinal cup to be flattened. Fine insect pins were used to hold all the petals of the retinal cup flat, taking care to avoid puncturing the retina by pinning through the remaining iris at the periphery. Pointed tweezers were used to gently split the retina and sclera, separating the connection of the peripheral edge of the retina to the underlying epithelial layer, repeating the process until all the edges of the retina were free. The free-floating retina was then gently transferred onto a glass slide, retinal cup-side up, and flattened using closed tweezers or a fine brush to unfold the petals. A small square of filter paper was then placed over the flat retina to adhere it to the retina. The retina and filter paper were then transferred back into 4% PFA for a secondary 15-minute fixation. The retina on filter paper was then transferred into a fresh tube of PBS, and the retina was gently pushed off the filter paper so that it was free-floating. The retina was then rinsed five minutes before starting the whole-mount immuno-staining protocol or it was stored in PBS at 4°C overnight for immunostaining the next day.

### Immunostaining

For immunocytochemical staining of optic nerves and optic tracts, a blocking solution of 5% normal goat serum (Jackson ImmunoResearch), 0.5% Triton X-100 (Sigma-Aldrich), and 0.5% bovine serum albumin (BSA; Sigma-Aldrich) in PBS was applied for 2 h. Primary antibodies were diluted in the blocking solution and incubated overnight at 4°C. Slides were washed 3 times for 10 minutes with PBS and 0.5% Triton X-100 before secondary antibodies were applied for 2 h at room temperature at 1:1000 dilution in PBS, 0.5% Triton X-100, and 0.5% BSA. Secondary antibodies were centrifuged at 13,000 rpm for 10 minutes at 4°C before application. Slides were washed 3 times for 10 minutes with PBS and 0.5% Triton X-100 and finally PBS alone for 10 minutes before being sealed with Prolong Diamond Antifade Mountant with DAPI (catalog #P36962, Thermo Fisher Scientific).

Retinas, isolated and fixed as described above, were rinsed with PBS for five minutes before adding blocking solution consisting of 5% normal goat serum (Jackson ImmunoResearch), 0.5% Triton X-100 (Sigma-Aldrich), and 0.5% bovine serum albumin (BSA; Sigma-Aldrich) in PBS, which was applied for at least one hour at room temperature with gentle shaking. Primary antibodies were diluted in the blocking solution and retinas were incubated for 48 hours at 4°C. Slides were washed 4 times for 10 minutes with PBS and 0.5% Triton X-100 before secondary antibodies were applied overnight at 4°C at 1:1000 dilution in PBS, 0.5% Triton X-100, and 0.5% BSA. Secondary antibodies were centrifuged at 13,000 rpm for 20 minutes at 4°C before application. A 1:1000 dilution of DAPI was added at the same time as secondary antibodies. Slides were washed 4 times for 15 minutes with PBS and 0.5% Triton X-100. Retinas were gently transferred from a tube to a slide, carefully avoiding pinching tissue. A drop of Prolong Diamond Antifade Mountant with DAPI (catalog #P36962, Thermo Fisher Scientific) was added to the retina on the slide.

### Confocal Microscopy

Images were acquired by confocal laser scanning microscope using Olympus Fluoview software (model Fluoview FV3000, Olympus) using an Olympus PlanApo N 60x/1.42 n.a. oil objective lens. All images were coded and analyzed without the knowledge of the experimental condition by the investigator making the measurements.

Nodes of Ranvier were immunohistochemically stained for Contactin Associated Protein 1 (Caspr1), a paranodal region-associated cell adhesion molecule expressed on the flanks of nodes of Ranvier, and by sodium channel Scn8a (Nav 1.6), which is concentrated within the nodal gap. Visualization of retinal tissue involved immunohistochemical staining for RNA-binding protein with multiple splicing (RBPMS) for retinal ganglion cells (RGC), and red fluorescent protein (RFP) to visualize viral transfection of RGCs, with DAPI staining to visualize cell nuclei.

### Electron Microscopy

Mice were deeply anesthetized and perfused with 2.5% gluteraldehyde and 4% PFA in 0.1 M sodium cacodylate, pH 7.4. Optic nerves were immediately removed from animals, then post-fixed in the same solution for 2 hr. Samples were post-fixed for 1 hr in 2% osmium tetroxide in 0.1M sodium cacodylate, pH 7.4. Samples were washed twice in 0.05 M sodium acetate, pH 5.0, and stained with 2% uranyl acetate in 0.05 M sodium acetate, pH 5.0, for 2 hr at 4°C. Samples were then dehydrated through a graded series of ethanol solutions followed by 100% acetone, and infiltrated with Epon 812 (Electron Microscopy Sciences, Hatfields, PA). Embedded samples were polymerized in an oven set at 60°C. Ultrathin sections (90 nm) were cut with a diamond knife on a Leica EM UC7 Ultramicrotome. Sections were placed on 200 mesh copper grids and post-stained with, uranyl acetate and lead citrate and examined by transmission electron microscopy on a JEOL-1400 Transmission Electron Microscope operating at 80 kV and with an AMT BioSprint-29 megapixel camera with 9.5 micron square pixels.

All images were coded and analyzed without the knowledge of the experimental condition by the investigator making the measurements. The diameter of all axons cut in cross section was measured, both for the axon itself and the fiber diameter including the myelin sheath. Transected axons are rarely perfect circles, so both the major and minor axes of each cross sectioned axon were measured and averaged to calculate the g-ratio as the axon diameter divided by the fiber diameter. Measurements of axon diameter, fiber diameter, number of axons, and g-ratios of all axons in the field of view were averaged and used for statistical testing, comparing between experimental and control conditions using unpaired two-tailed Student’s *t*-tests. This design avoids pseudo-replication that would result from using each axon as the sample size in statistical test calculations.

### Image Acquisition and Nodal Gap Measurement

In both the monocular deprivation and monocular inhibition paradigms, an Olympus PlanApo N 60x oil microscope objective (60x/1.42 n.a. Oil; UIS2 BFP1; ∞/0.17/FN26.5) with Olympus IMMOIL-F30CC immersion oil type-f was used at a 4x zoom. Olympus Fluoview software was used for image acquisition and analysis. A z-series of ten sections of 0.2 µm intervals was taken for optic nerve samples for the monocular inhibition paradigm, and a z-series of 30 sections at 0.2 µm intervals was taken for optic tract analysis, for both the monocular deprivation and monocular inhibition experiments with the opposite eye of the same animal used as a control. Thicker samples were required in the optic tract because axons are less highly compacted than in the optic nerve.

Ten regions of interest along the full length of the optic nerve or tract were imaged at random locations. A single-plane maximum projection of the z-series stack was created using Olympus Fluoview and NIH ImageJ software. In samples with virus-infected RFP+ axons, a single-plane maximum projection was not created, and each stack was reviewed one image at a time to ensure co-localization of RFP+ axons and Caspr1. Nodal gap lengths were measured using NIH ImageJ software by using the line measurement tool to measure from the edge of one Caspr1-labeled paranode to the opposing paranode edge delimiting the nodal gap. The mean nodal gap for all nodes of Ranvier in the field of view was calculated and used for statistical analysis, rather than using each node of Ranvier as n = 1. Thirty Z-plane stacks of 53 X 53 µm^2^ confocal fields of view collected at 0.2 µm intervals represent a sampled depth of 6.6 µm, and a volume of 0.185 mm^3^ considering the approximately 0.8 µm depth of a single confocal image plane. All samples were coded and measured without knowledge of the experimental conditions. A total of 35,560 nodes of Ranvier from a total of 6,600 images were used for this study.

In the monocular inhibition paradigm, retinal ganglion cell (RGC) numbers were counted in the retinas of postnatal day 100 (P100) animals to ensure that the virus was not significantly altering the number of RGCs and to determine the proportion of cells overexpressing Kir2.1. Flat-mount retinas were stained with anti-RNA-binding protein with multiple splicing (RBPMS) antibodies and DAPI to visualize total RGC numbers and to confirm that a signal in the image is a cell. If a cell had both the RBPMS signal and was RFP+, a RGC had been successfully infected with the virus. Flat-mount retina images were captured by a confocal laser scanning microscope with an Olympus UPlanSApo 20x microscope objective (20x/0.75; UIS2; ∞/0.17/FN26.5) at a 1x zoom using the Olympus Fluoview software. A z-stack of 30 sections at 0.2 µm intervals was collected at five random points in the retinas. A single-plane maximum projection was formed for each flat-mount retina image and the point tool was used to count individual cells and record whether they are RFP+. A total of 29,885 RGCs were counted from the retinas of N=3 Vglut2-Cre animals of both sexes.

## Electrophysiology

### Pattern Electroretinogram (pERG) and Pattern Visual Evoked Potential (pVEP)

After 30 days of MD, sutures were removed from the deprived eye under dim red light to preserve dark adaptation in mice, which lack L-opsin photoreceptors and thus cannot detect red wavelengths (Greenwald et al., 2014). Mice were anesthetized using Ketamine (0.1 mg/g) and Xylazine (10 mg/kg), and they were placed on the testing platform for recording visually evoked electrophysiological responses (Diagnosys Celeris apparatus), maintaining body temperature of 37°C. Mice were treated prior to pVEP and pERG analysis with Proparacaine Hydrochloride Ophthalmic Solution (USP 0.5%:: NDC 61314-016-01) to relieve discomfort of the probe and miniature LCD display on the eye. Cyclopentolate Hydrochloride Ophthalmic Solution (USP 1%: NDC 61314-396-03) and Phenylephrine Hydrochloride Ophthalmic Solution (USP 10%: NDC 42702-103-05) were administered to dilate the pupil. All ophthalmic solutions and drugs used were acquired through the NIH Veterinary Department.

Visual stimulation using bars of 50cd/m^2^ intensity was delivered by a miniature LCD display placed directly on the corneas. Horizontal bar patterning was presented to the animals, with 4 bars in the image sweeping the retina at 2 cycles per degree. Two sets of 300 pERG and pVEP runs were recorded and averaged. Each eye was tested sequentially. The ground lead was inserted into the hind flank of the mice, and the recording lead for pVEP was inserted subdermally on the skull above the visual cortex at the midline. The pattern stimulator was placed on the eye being tested, and an unused full-field stimulator was placed on the other eye as an electrical control. Stimulation information was recorded using Diagnosys Espion V6 software. The animals’ eyes were treated with TAOO to prevent drying.

### Statistical Analysis

All samples were coded and analyzed by a different investigator without knowledge of the experimental condition. Statistical analysis was performed using GraphPad Prism (v10.0). Normality was assessed using the Shapiro-Wilk or Kolmogorov-Smirnov test for the data reported. All data were reported as mean ± SEM, and two-sided statistical comparison was utilized (unless otherwise noted) and considered significant at p < 0.05. To assess whether differences in pVEP latency between the two eyes of individual animals differed significantly from the expected ratio of 1, a one-tailed t-test was used. This approach eliminated variation in electrophysiological data among different animals by using each animal as its control.

Pseudo-replication was avoided in nodal gap length and ultrastructural data by averaging the values of all measured structures within a confocal z-stack or EM field, with each field being treated as an independent replicate. Unpaired two-tailed Student’s *t*-tests were used to compare the nodal gap lengths on axons in the same tissue conveying action potentials driven by normal vision vs. axons with impulse activity perturbed by MD or monocular inhibition (MI). Nodal densities were calculated by taking the total number of nodes per field and comparing the respective optic nerves and optic tracts. The ratio of latency to peak pVEP from the two eyes was analyzed using one-sample Student’s *t*-tests for deviation from an expected ratio of 1 (equal latency from both eyes). The use of each animal as its own control reduces experimental variance among animals inherent in electrophysiological recording due to animal size and electrode placement, and physiological variance in response latency among animals.

## Results

### Plasticity of Nodes of Ranvier in Optic Tracts Following MD

We examined the effect of MD on Nodes of Ranvier in optic nerves and optic tracts using confocal microscopy after immunohistochemical identification of Contactin-associated protein 1 (Caspr1) in the paranodal region, and sodium channels Nav1.6 in the nodal gap (**Fig. 1A**). Intraocular injection of red fluorescent cholera toxin B conjugated to AlexaFluor-555 (CTB-555) into the eye with eyelid suture was used to identify nodes on axons from the sutured eye (**Fig. 1B**). CTB-555 accumulates in the paranodal region.

**Fig. 1.**
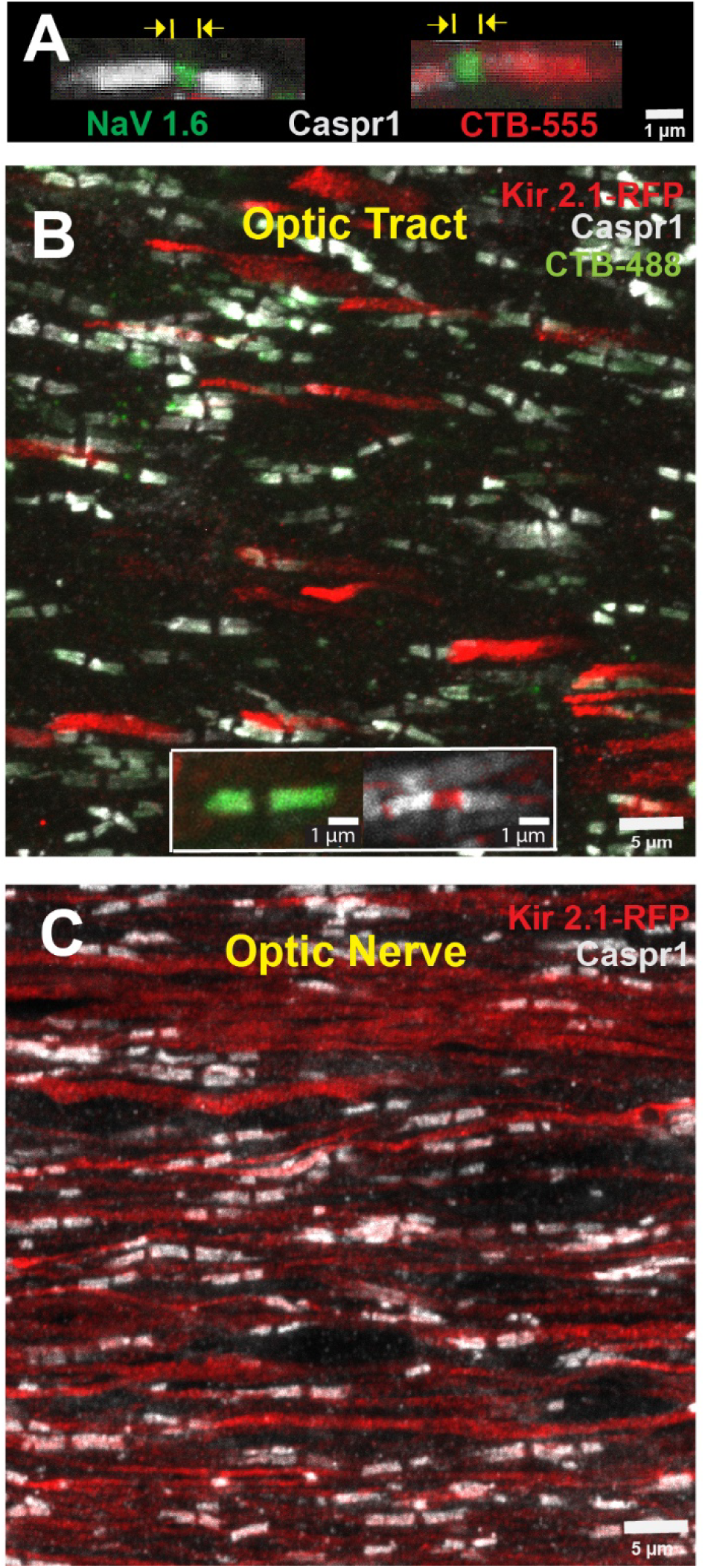
Nodes of Ranvier in mouse optic nerves and tracts after monocular deprivation and monocular inhibition. **(A)** Nodes of Ranvier (NOR) were identified by using antibodies for Contactin-associated protein 1 (Caspr1) to label paranodes (grey), and sodium channels SCN8A (NaV1.6) within the nodal gap (green). Intravitreal injection of AlexaFluor 555-conjugated cholera toxin subunit B (CTB-555), which accumulates in the paranodal regions, tags nodes on axons originating from the sutured eye (red). Nodal gap lengths were measured from paranode to paranode for all nodes in the field of view. For monocular inhibition (MI), to inhibit action potential firing in a subset of axons in optic tract **(B)** and optic nerve **(C)**, Kir2.1 potassium channel was transfected into retinal ganglion axons to hyperpolarize them by intraocular injection of AAV2-EF1a-FLEX-Kir2.1-T2A-td Tomato. Transfected axons were identified by the red tdTomato fluorescence (Kir 2.1 RFP). Axons originating from the other eye were identified by injection of CTB 488 into that eye, which accumulates at paranodal junctions. High-magnification insert is shown in B.

The results showed that NOR were significantly shorter on optic tract axons from the deprived eye compared to axons from the unsutured eye in the optic tract ipsilateral to the unsutured eye (p < 0.003, t-test_(58 df)_ = 3.12) (**Fig. 2**). An opposite effect was apparent in the contralateral tract, with a strong trend for NOR on axons from the deprived eye to be longer than those from the open eye. However, the statistical probability of the difference was below the p < 0.05 threshold criterion (p = 0.11, t-test_(58 df)_ = 1.62) (**Fig. 2**).

**Fig. 2.**
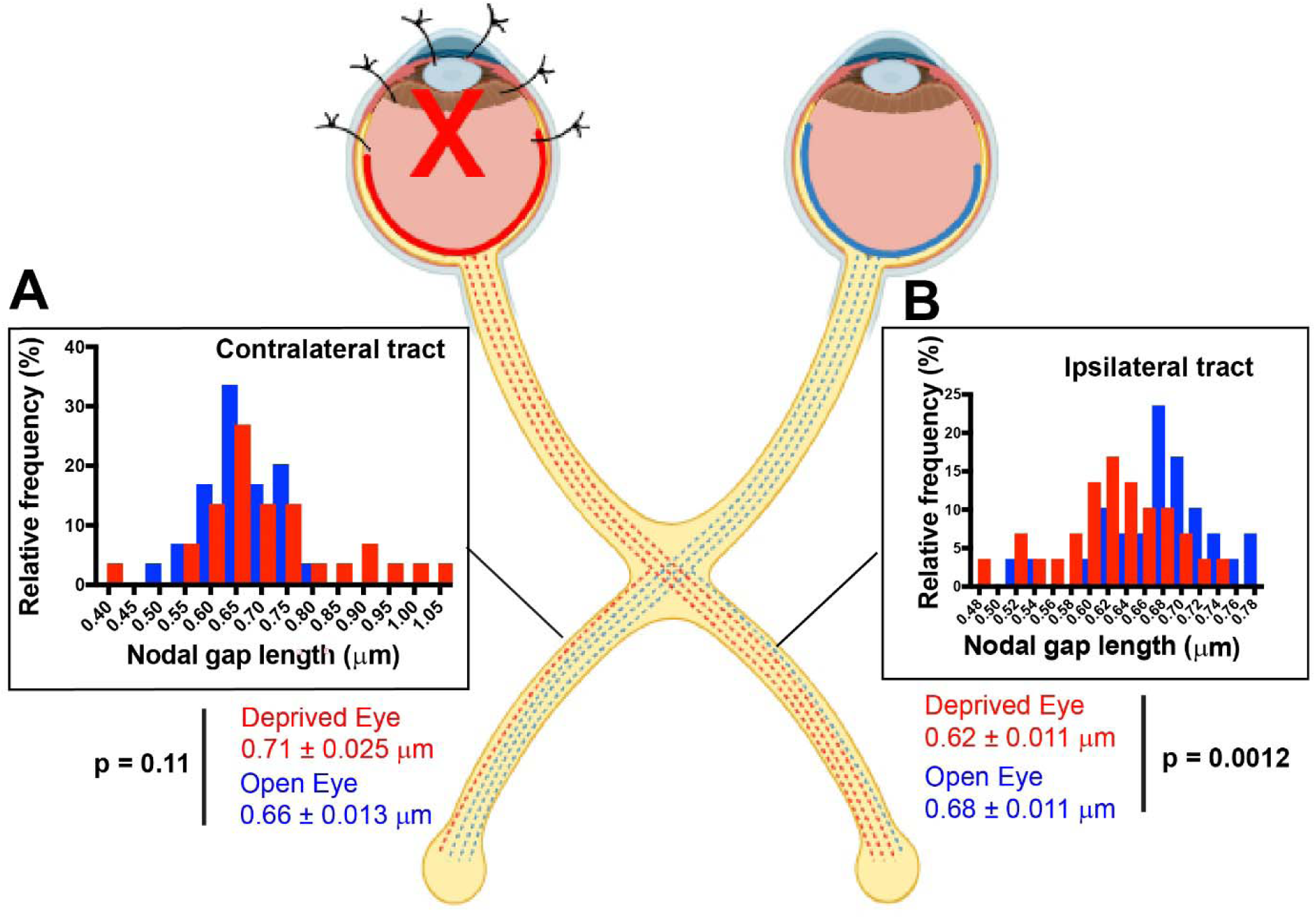
Monocular deprivation by eyelid suture of adult mice alters the length of nodes of Ranvier in optic tracts. Nodal gaps were significantly shorter on axons from the deprived eye (red) compared to axons from the uninhibited eye (blue) in the optic tract ipsilateral to the open eye (p = 0.0012, t-test_(df= 58)_ = 3.403). In the contralateral optic tract, there was a strong trend for nodes to be longer on axons from the deprived eye (red) relative to axons from the open eye (blue) in the same tract (p = 0.11, t-test_(df= 58)_ = 1.618), but the differences failed to reach statistical significance at the p < 0.05 probability level. Nodal size frequency histograms are shown for nodal gap lengths measured on axons averaged by field from the sutured eye (red) and unsutured eye (blue) for the contralateral (**A**) and ipsilateral optic tracts (**B**). Shorter nodal gaps are known to increase conduction velocity (8).

Different effects of NOR plasticity in the ipsilateral and contralateral optic tracts parallels the different effects found in ipsilateral and contralateral visual cortex of adult mice in ocular dominance studies of synaptic plasticity (Sawtell et al., 2003). MD by eyelid suture for 12 days in mice between the ages of P60-90 leads to persistent NMDA receptor-dependent enhancement of visual evoked potential (VEP) responses from the open eye in the visual cortex ipsilateral to the open eye, but little or no change in deprived eye responses in that cortex (Frenkel and Bear, 2004; Sawtell et al., 2003). This is attributed to the fact that 90 percent of axons decussate, resulting in much lower levels of impulse activity in the cortex contralateral to the sutured eye. This is believed to induce a process of metaplasticity, in which the threshold and direction of synaptic strength modification depend on the prior level of ongoing postsynaptic activity (Abraham and Bear 1996; Cooper and Bear 2012). Metaplasticity does not predominate in visual cortex contralateral to the open eye, because afferent activity is reduced in only 10 percent of the axons. Here the effects are variable, depending on the interplay between STDP and metaplasticity, which vary with experimental parameters; notably postnatal age and the duration of visual deprivation (Sawtell et al., 2003; Frenkel and Bear 2004). Synaptic competition predominates in young mice, thereby decreasing responses to the sutured eye relative to the open eye in both cortices; however, this process abates in older mice after the close of the critical period. The possibility of changes in myelin in the optic tract axons was not considered previously.

The finding of significantly shorter NOR on axons from the MD eye in ipsilateral optic tract reveals a form of NOR plasticity and raises the question of whether neural transmission speed is altered in axons from the visually deprived eye. This was tested by determining the latency of visually-evoked potentials in the visual cortex from the open eye and the eye that had been sutured for 30 days.

### Effect of MD on Spike Time Arrival in Visual Cortex

Mathematical modeling and electrophysiological measurements in mouse optic nerve show that lengthening the nodal gap by 0.14 µm results in approximately 20% slower action potential transmission speed, and 6.7 ms increased latency of spike time arrival at the visual cortex (Dutta et al., 2018). This difference in spike time arrival falls within the temporal window of spike time arrival synchrony with respect to postsynaptic action potential firing to induce STDP plasticity (Dan and Poo, 2006; Mu and Poo, 2006; Andrade-Talavera et al., 2023). A nodal gap length increase of this magnitude is functionally significant, with a reduction in visual acuity of about 0.01 cycles per degree (Dutta et al., 2018). Also, in wild-type rat optic nerve and cortex, NOR gap lengths vary over a 4.4 and 8.7-fold range respectively in each tissue, and mathematical modeling predicts that these nodal length differences will alter conduction speed by ∼20% (Arancibia-Carcamo et al., 2017). The significantly shorter NOR gap length on axons from the deprived eye in the optic tract ipsilateral to the open eye would predict an increase in conduction velocity in these axons.

To determine whether MD alters spike time arrival in the visual cortex, the latency to peak visually evoked potential using patterned visual stimulation (pVEP) was determined by inserting a subcutaneous recording electrode against the skull above the visual cortex at the midline. After 30 days of MD, sutures were removed from the deprived eye under dim red light to preserve dark adaptation in mice for pVEP measurements. The measurements showed that the latency to peak pVEP was shorter in every animal from its deprived eye compared to its unsutured eye, with a mean decrease in latency of 13.9% from the deprived eye of mice that had undergone 30 days MD (p = 0.004, t-test_(3 df)_ = 8.040) (**Fig. 3**). Comparing latencies of VEP responses from both eyes of the same animal eliminates inter-animal variability in pVEP response latencies. As expected, in mice that did not experience MD, the ratio of latencies to peak pVEP response from the two eyes using each animal as its own control was not significantly different from a ratio of 1 (p = 0.19, one-tailed t-test _(4 df)_ = 1.575) (**Fig. 3B**). In addition, as expected (Oner et al., 2004), the mean amplitude of pVEP evoked by the deprived eye (-5057 ± 1750 nV) was about half that of the nondeprived eye (-9535 ± 624.8 nV), (p = 0.05, t-test_(6 df)_ = 2.410) (**Fig. 3C**). Thus plasticity of NOR gap length after MD in adult mice changes conduction velocity in a manner that would disrupt the synchrony of synaptic input from deprived-eye axons with respect to action potential firing in the post synaptic binocular neuron, reducing the amplitude of those epsps.

**Fig. 3.**
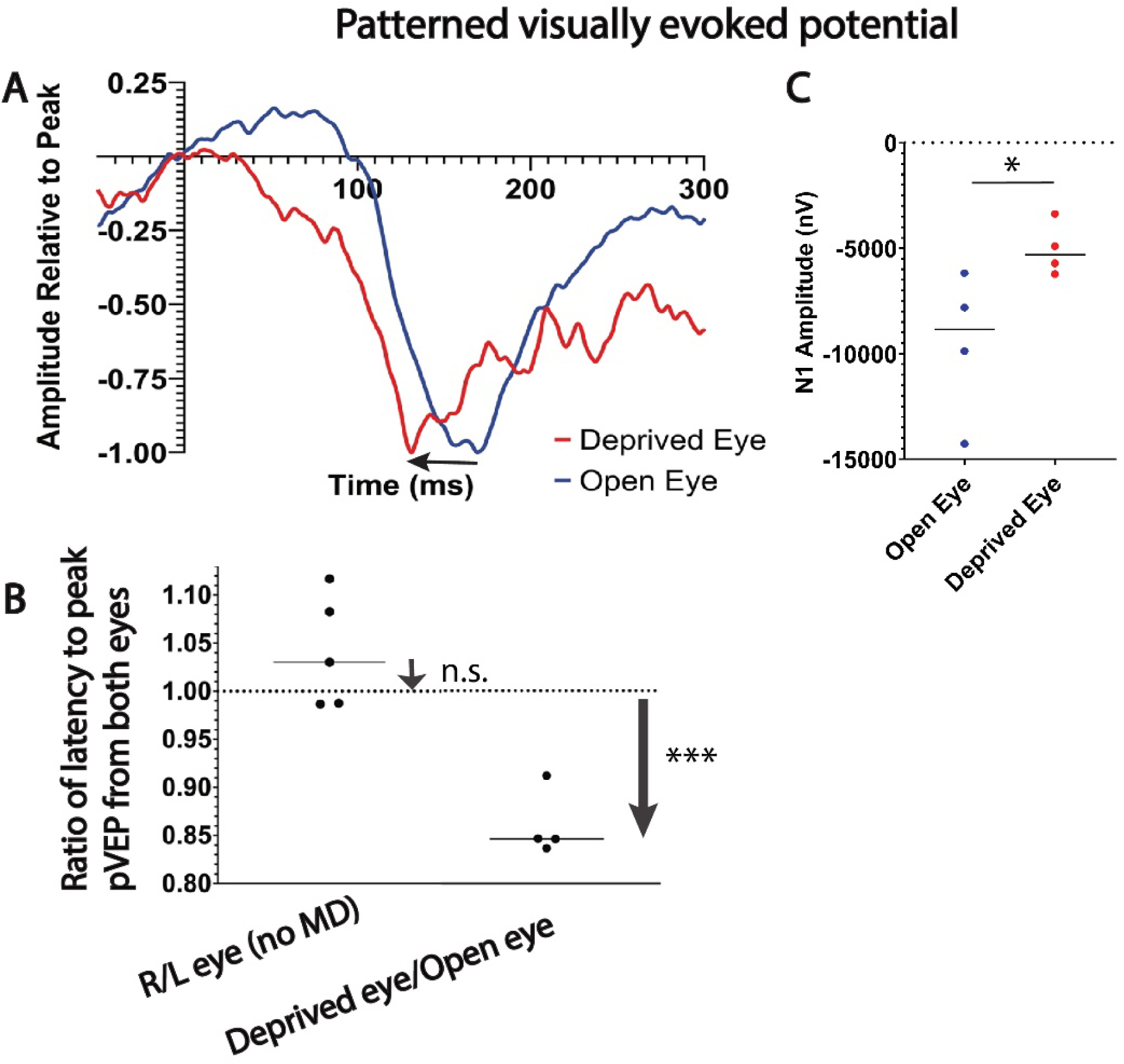
Monocular deprivation reduces the latency and amplitude of visually evoked potentials in visual cortex from the deprived eye. (A) Representative patterned visually evoked potential waveforms from the unsutured eye (blue trace) and previously sutured eye (red trace) of an adult mouse, showing the shortened latency to peak pVEP (arrow) from the eye that had been deprived of vision for 30 days. (B) The ratio of latency to peak pVEP from the two eyes was not significantly different from an expected ratio of 1 in mice with normal visual experience (ratio of 1.04 right eye/left eye ± 0.06, p = 0.19, one-tailed t_(4 df)_ = 1.575), but was significantly reduced from the deprived eye (ratio of 0.86 ± 0.03 MD/open eye, t_(3 df)_ = 8.040; p = 0.004), consistent with faster conduction velocity in MD axons. (C) The amplitude of pVEP responses were smaller from the eye that had been sutured (p = 0.05, two-sample t_(6 df)_ = 2.410). *p < 0.05; *** p < 0.0004, n.s. non-significant.

### Oligodendrocytes as Mediators of Action Potential Synchrony

As currently understood, metaplasticity and homeostatic plasticity depend on the level of postsynaptic activity, but this information is not directly accessible to oligodendrocytes. Recently a theory of oligodendrocyte mediated plasticity (OMP) has been proposed to address this problem (Pajevic et al., 2023). Action potentials trigger localized intracellular calcium responses in oligodendrocyte processes at sites of axon contact through the vesicular release of glutamate from axons activating mGluR and NMDA receptors on oligodendrocytes (Wake et al., 2011). This intercellular signaling stimulates activity-dependent myelination by increasing local translation of myelin basic protein (Wake et al., 2011). Thus, action potentials in axons are detected at points of contact between the oligodendrocyte’s individual processes associated with different axons.

Mathematical modeling indicates that if the timing of spikes arriving among the population of axons that are myelinated by a single oligodendrocyte initiate changes in myelin on each axon to minimize temporal delays of action potentials among the local population of axons that the oligodendrocyte contacts, then the synchrony of spike time arrival at the axon terminal is greatly improved (Pajevic et al., 2023). The modeling further shows that non-coherent firing among axons does not induce OMP. In nerves and tracts with mixed populations of axons having different firing patterns, groups of axons are sorted by their coherent activity for alterations in myelination.

To test the hypothesis that NOR plasticity is driven by differences in activity among individual axons in the visual pathway, rather than a tissue-level effect, a mixture of functionally active and inactive axons was imposed in the optic nerve to mimic the situation in optic tract. Previous research shows that 30 days binocular deprivation does not alter nodal gap width significantly on axons in the optic nerve (Santos et al., 2022), but there is no binocular information in the optic nerve. To introduce differences in action potential activity among axons in the optic nerve, neural impulses were suppressed in a subset of axons in optic nerve by transfecting inwardly rectifying potassium channel (Kir2.1) selectively into retinal ganglion axons of one eye by intraocular injection of AAV2-EF1a-FLEX-Kir2.1-T2A-tdTomato (Cue et al., 2014). Histological analysis of retinas showed potassium channel Kir2.1 expression in approximately 41.3% of RGCs (**Fig. 4 A, B**). Electrophysiological recording of pattern electroretinograms (pERGs) confirmed that retinal responses were significantly attenuated through the course of the experiment by the overexpression of Kir2.1 to inhibit action potential firing in RGCs (p = 0.0021; t-test_(10 df)_ = 4.112) (**Fig. 4 C, D**). NOR on axons that were successfully transfected were identified by the red fluorescence of tdTomato co-expressed by the virus.

**Fig. 4.**
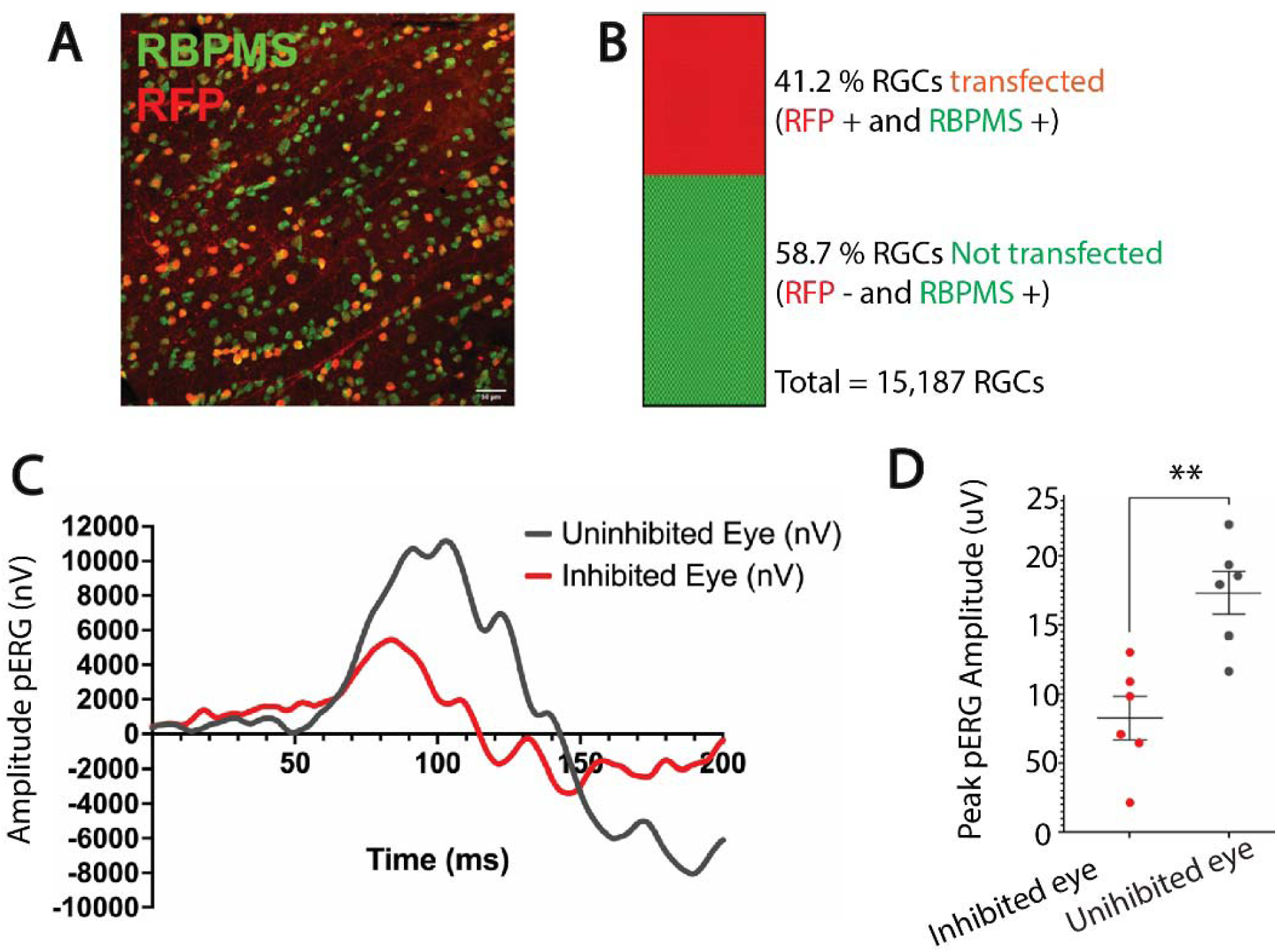
Overexpression of the inwardly rectifying potassium channel 2.1 reduces the amplitude of pattern electroretinogram responses (pERG). **(A)** Representative confocal microscope field of retinal ganglion cells in the monocularly inhibited retina of adult mice measured at postnatal day 100. Retinal ganglion cells were labeled with antibodies for RNA-binding protein with multiple splicing (RBPMS) (green). Red fluorescent protein (RFP) (red) indicates retinal ganglion cells transfected with Kir2.1. (**B**) Retinas from the eyes injected with AAV2-EF1a-FLEX-Kir2.1-T2A-tdTomato had an average of 41% of the RGCs transfected with Kir2.1 to inhibit action potential firing (red), and 58% were not transfected (green) N = 3 mice. (**C**) Representative electroretinogram response to patterned visual stimulation showing greatly reduced responses after inhibiting action potentials in approximately half of the RGCs of one eye (red), compared to responses in the other eye of the same mouse that was not transfected (black). (**D**) Mean peak pERG amplitudes were significantly reduced in retinas after Kir2.1 transfection compared to uninhibited retinas (4123 ± 786.9 nV vs. 8672 ± 777.6 nV; p = 0.0021 t-test_(10 df)_ = 4.112). * p < 0.05; ** p < 0.005. There was no significant difference in the total number of retinal ganglion cells between the monocularly inhibited eye and the uninhibited eye (p = 0.6, t-test_(28 df)_ = 0.4868).

The results using Kir2.1 transfection into RGCs showed that when oligodendrocytes in optic nerve are provided access to a mixed population of axons with either visually evoked activity or inhibited action potential activity, activity-dependent differences in NOR length resulted which were axon-specific (**Fig. 5**). Nodal gap length was increased selectively on optic nerve axons in which action potentials were inhibited by expressing Kir2.1 (p = 0.0001, t-test _(118 df)_ = 6.41).

**Fig. 5.**
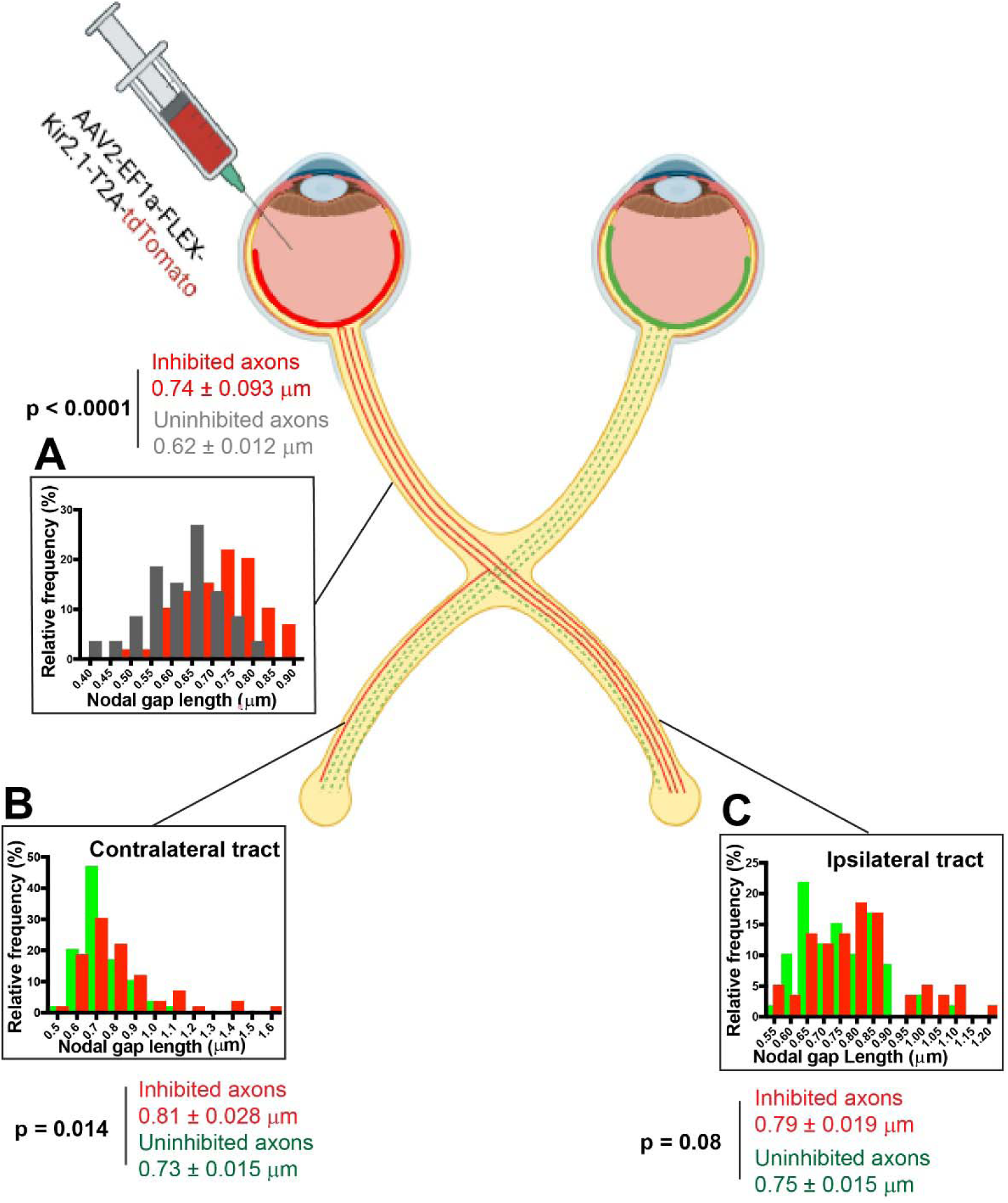
Inhibiting action potential firing in a subset of axons from one eye, reveals an activity-dependent axon-specific mechanism of nodal plasticity. (**A**) In the optic nerve, nodes of Ranvier were significantly larger on axons transfected with Kir2.1 to inhibit action potentials (red) compared to untransfected axons (grey) in the same nerve (p = 0.0001, t-test_(118 df)_ = 6.41). (**B**) In the optic tract contralateral to the untransfected eye, nodes of Ranvier were significantly larger on axons with action potentials inhibited (red) compared to untransfected axons (green) in the same tract (p = 0.014, t_(118 df)_ = 2.48). (**C**) A similar trend was observed in the optic tract ipsilateral to the open eye, with larger nodes of Ranvier on axons with action potentials inhibited (red) compared to untransfected axons (green) in the same nerve, but the differences did not reach the 0.05 level for statistical significance (p = 0.08, t-test(_118 df)_ = 1.76).

Monocular inhibition (MI) of action potential firing also resulted in lengthening NOR gaps on axons with inhibited action potential firing in the optic tract contralateral to the open eye (p = 0.014, t-test _(118 df)_ = 2.48) (**Fig 5**). A similar effect was apparent in the ipsilateral optic tract (p = 0.08, t-test _(118 df)_ = 1.76), but the differences failed to reach the p < 0.05 level of statistical significance.

Together these results reveal that in addition to modification of synaptic strength in the lateral geniculate nucleus and visual cortex, MI and MD induce axon-specific activity-dependent changes in NOR gap length that will affect the speed of impulse transmission to the CNS. This is consistent with an axon-specific mechanism of NOR plasticity that is dependent on differential levels of neural impulse activity in individual axons, rather than a tissue-level effect, as for example could result from activity-dependent release of neurotrophic factors.

### Ultrastructural Differences Following MI

In addition to changes in NOR gap length, several other factors contribute to spike time arrival in visual cortex, including synaptic delays, myelin thickness, axon diameter, and ion channel composition in the nodes. Previous studies have shown no differences in axon diameter or myelin thickness in optic nerve following visual deprivation (Etxeberria et al., 2016). Possible ultrastructural differences in optic nerve axons after MI were analyzed in the present experiments by transmission electron microscopy.

At the ultrastructural level, axon morphology and myelin appeared normal in optic nerves after MI, with no evidence of axon degeneration (**Fig. 6 A-C**). Quantitative analysis revealed no significant difference in mean myelin thickness as expressed by g-ratio in the two optic nerves of the same mice after MI inhibition (g-ratio = 0.80 ± 0.0042 vs 0.79 ± 00038, inhibited nerve vs uninhibited nerve (p = 0.20, t-test _(57 df)_ = 1.295) (Fig. 6 D).

**Figure 6.**
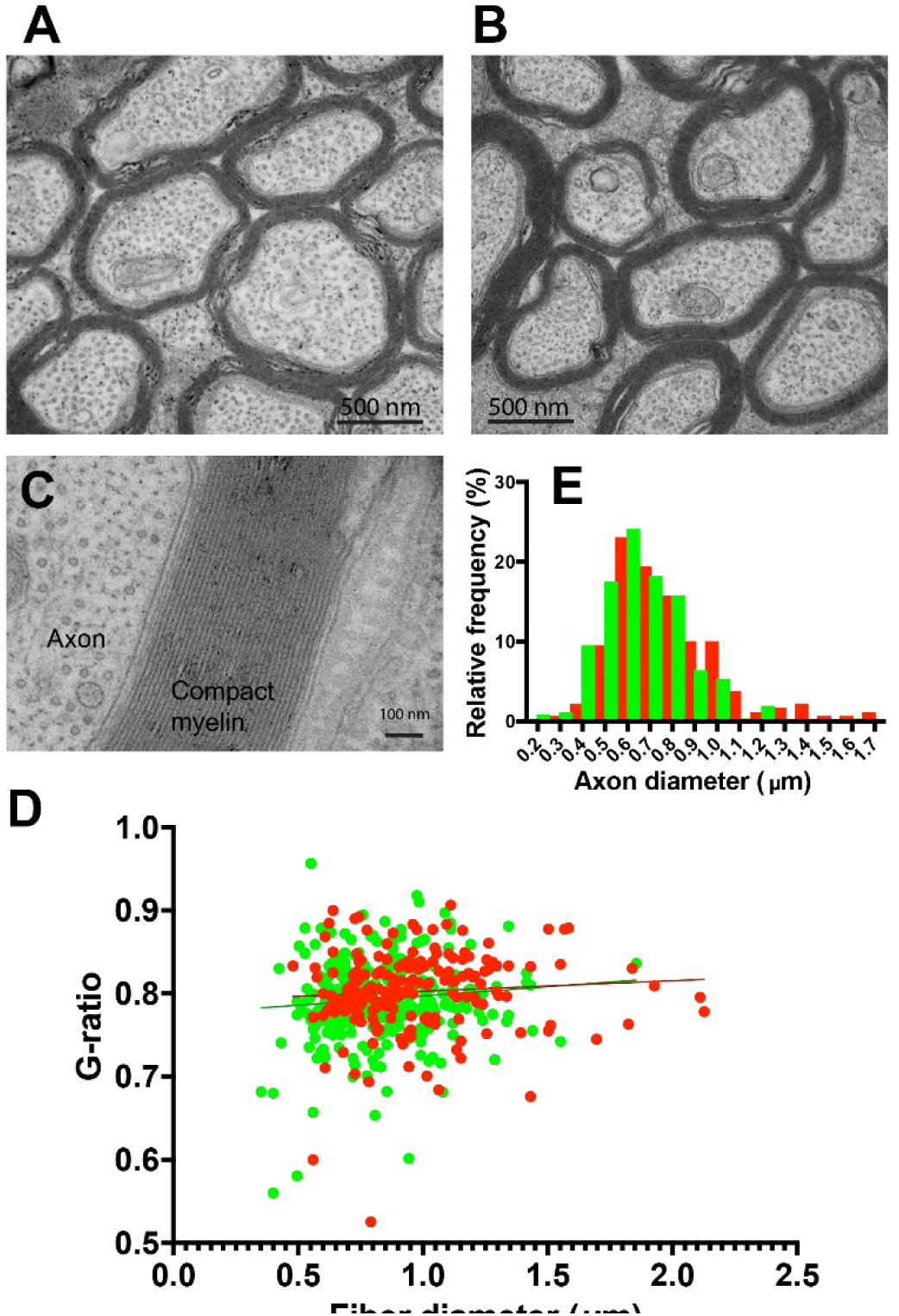
Ultrastructure of axons in optic nerve after MI by viral transfection of Kir2.1 to inhibit action potential firing. Normal appearing ultrastructure of axons in the optic nerve from the transfected eyes. (**A**), compared to the untransfected eyes (**B**). (**C**) High magnification showing normal compaction of myelin after MI. (D) There was no significant difference in myelin thickness (g-ratio) in transfected optic nerve axons after MI (red) compared to axons in the untransfected eyes (green), mean 0.80 ± 0.0041 vs 0.79 ± 0.0038 transfected (red) vs untransfected (green) eyes, (p = 0.20, t-test_(57 df)_ = 1.29). The plot shows 192 axons from the transfected optic nerve and 288 axons from the untransfected nerve. (E) The mean caliber of axons was slightly larger in the MI condition 0.79 ± 0.19 µm vs 0.68 ± 0.18 µm, transfected (red) vs untransfected eye (green), (p = 0.0001, t-test_(55 df)_ = 4.136).

There were, however, slightly fewer axons in the optic nerve transfected with Kir 2.1 (16.5 ±0.75 vs 19.5 ± 0.6 axons/29.7 μm^2^, p < 0.003, t-test _(57 df)_ = 3.07), and consequently, fewer NOR in the optic nerves transfected with Kir 2.1 (80.4 ± 2.59 vs. 95.9 ± 3.18 nodes/7303μm^3^, p < 0.0002, t-test_(118 DF)_ = 3.80). Since the control eye was injected with CTB-488 tracer, the difference in the number of axons is not likely caused by the injection, but rather a result of Kir2.1 expression.

This suggests that in addition to activity-dependent competition affecting NOR gap length on axons from the retinas, there is also a deprivation-dependent process of axonal atrophy after prolonged inhibition of action potentials in optic nerves of adult mice. There is little prior research on possible loss of axons after prolonged action potential inhibition in adults, but during development, activity-dependent access to trophic factors and activation of apoptotic pathways eliminates ineffectual axons and neurons (Linden, 1994 for review).

A small but significant increase in mean axon caliber was also evident by electron microscopy in optic nerve axons from the eye experiencing MI (0.79 ± 0.019 vs 0.68 ±0.18 µm, inhibited nerve vs. uninhibited nerve, p = 0.0001, t-test _(55 df)_ = 4.13) (**Fig. 6E**). There is no relation, however, between nodal gap length and axon caliber in optic nerve (Arancibia-Carcamo et al., 2017). In theory, increased axon caliber would tend to increase conduction velocity in transfected axons, but the question is largely moot, as action potential firing is inhibited in these axons. A limitation of the electron microscopy data is that unlike NOR measurements by confocal microscopy, where individual axons from each eye are labeled with different fluorescent indicators, the ultrastructural data in optic nerve from the MI eye represent mean values comprised of a mixture of axons with action potentials inhibited by Kir2.1 transfection, and axons with normal visually evoked activity.

## Discussion

The implicit assumption in studies of ocular dominance plasticity that the speed of impulse transmission is the same from both eyes and is not altered by visual experience is not supported by these experimental results. The data show that MD alters the latency of spike time arrival in visual cortex, with shorter latency visually evoked potentials from the eye deprived of vision by eyelid suture for 30 days in adulthood. This is accompanied by morphological plasticity of nodes of Ranvier, resulting in shorter nodal gaps on axons from the deprived eye in optic tract ipsilateral to the open eye.

An axon-specific mechanism of NOR plasticity is demonstrated by the significantly different NOR gap lengths on axons from the eye deprived of vision by eyelid suture compared to axons from the non-deprived eye in optic tracts. The lack of changes in NOR length in optic nerves of mice deprived of binocular vision for 30 days (Santos et al., 2022; Santos et al., 2024) suggests a plasticity mechanism dependent upon differences in activity among individual axons. Confirming this experimentally, when differences in activity among axons in optic nerve were introduced by inhibiting action potentials in a subset of axons by transfection of Kir2.1, nodal gap length was increased specifically on the inhibited axons. This supports a competitive activity-dependent mechanism of NOR plasticity as predicted by OMP theory (Pajevic et al., 2023).

Thus, upstream of synaptic plasticity detected in the CNS in ocular dominance studies, there is an axon-specific process of activity-dependent plasticity in the optic nerve and tract. NOR plasticity will affect action potential arrival time at binocular neurons, the coincidence of excitatory post-synaptic potentials (epsps) with respect to postsynaptic firing, temporal summation of postsynaptic potentials, and thereby the overall level of depolarization and action potential firing in the postsynaptic neurons. By STDP, these changes in impulse conduction speed would affect synaptic plasticity. In addition, changes in latency of spike time arrival will affect temporal summation of synaptic potentials in binocular neurons, potentially altering the waveform, latency, and amplitude of responses recorded in visual cortex. Since the relative amplitude of visually evoked potentials from the two eyes recorded in visual cortex is frequently used to monitor MD-induced ocular dominance shifts, the effects of spike time arrival on the waveform can be misinterpreted as exclusively reflecting changes in synaptic strength.

On average, there was a difference in latency in pVEP from the eye deprived of vision for 30 days and the open eye of 22.7 ms, which is in the range to reduce synaptic strength from its peak by STDP. Enlarged nodes on axons slows conduction velocity and by STDP this will also promote depression of synapses from these axons arriving after postsynaptic action potential firing.

The findings are compatible with the OMP model of activity-dependent myelin plasticity (Pajevic et al., 2023). The OMP model has three components. (1) a global, cell-body wide transient signal [G_(t)_], reflecting the combined activity of input from each of its processes in contact with an axon. (2) A local myelination factor [M_(t)_] released by each spike on a given axon, which is catalyzed by G_(t)_ (3) Two steady-state continuous processes of myelin addition (M), and myelin removal (τ). The widening of nodes when action potentials are suppressed may reflect the dynamic balance between myelin addition and removal, favoring removal when action potentials are inhibited.

In addition to the effects on NOR gap length from activity-dependent competition among axons, there was a small but significant decrease in the number of axons in optic nerves transfected with Kir2.1. This is consistent with a trophic or atrophy effect from suppressing neural activity for 30 days in adult mice occurring in tandem with the other mechanisms of plasticity.

Overall the findings suggest that oligodendrocytes serve to modify the transmission of information in axons in an activity-dependent manner analogous to the postsynaptic neuron in mediating activity-dependent synaptic competition (Malenka 1991) metaplasticity (Abraham and Bear, 1996; Cooke and Bear, 2010) and homeostatic plasticity (Ransom et al., 2012). Both synaptic and nodal plasticity emphasize the importance of synchrony of information transmission in activity-dependent plasticity, with synchrony of postsynaptic potentials assessed by the postsynaptic neuron and synchrony of action potentials in transmission to the target assessed by oligodendrocytes.

### Comparison with Other Studies

The effects of MD on NOR morphology have not been investigated previously. Prior research has shown that during early postnatal development in mice, monocular eyelid suture has several other effects, including increasing differentiation of oligodendrocytes and shortening the mean length of myelinated segments between NOR (Etxeberria et al., 2016). Curiously, the opposite effect on oligodendrogenesis is observed following optogenetic stimulation of mouse motor cortex, however (Gibson et al., 2014). The possible effects of MD on NOR gap length were not investigated in either study (Etxeberria et al., 2016; Gibson et al., 2014). In contrast to the axon-specific effects on NOR gap length found in the present studies on adult mice, the changes caused by MD during postnatal development and myelin formation were not specific to the axons from the deprived eye but also extended via an unknown mechanism to surrounding axons conveying normal vision (Etxeberria et al., 2016).

Comparing the present results on NOR plasticity during ocular dominance plasticity with similar studies of synaptic plasticity raises several questions. Direct comparisons are difficult, because the effects of visual deprivation on ocular dominance plasticity depend on the experimental parameters used, notably the species and age of the animal and the duration and means of visual disruption (Frenkel and Bear, 2004; Sawtell et al., 2003; Lunghi et al., 2015; Ransom et al, 2012). Experimental paradigms used to study ocular dominance plasticity vary widely, and none is identical to the procedure used here. Transfection of Kir2.1 was used in the present studies to provide an effective and long-lasting inhibition of action potential activity in RGC axons. Studies of inhibiting action potential activity on ocular dominance plasticity frequently use intraocular tetrodotoxin (TTX) injection to block voltage-dependent sodium channels, but this requires injections repeated daily to sustain inhibition (Frenkel and Bear 2004). Our preliminary experiments measuring pERG responses of retinal excitation showed that intraocular TTX injections inhibited retinal activity in mice for less than 24 hrs. Repeated injections to achieve 30 days of MI are not feasible, due to the trauma caused by daily injections. For this reason, prior studies of prolonged MI were limited to 3 daily TTX injections in adult mice for an experimental time-course of 5 days (Frenkel and Bear 2004). Moreover, Kir2.1 transfection achieves the experimental objective of suppressing the level of action potential firing in a subset of axons rather than blocking action potentials from retinas as in experiments using TTX. The effects of activity-dependent NOR plasticity revealed in the present experiments will need to be considered together with the well-established forms of synaptic plasticity in ocular dominance experiments.

The new findings also raise methodological issues that complicate interpretation of previous research, because ocular dominance studies frequently take the ratio of VEP amplitudes that are evoked by patterned visual stimulation of the two eyes to monitor ocular dominance shifts (Porciatti et al., 1999). This is a reliable indicator, but difficult to interpret at a cellular level. The changes in latency of spike time arrival of action potentials from the deprived eye will affect temporal summation and the waveform shape and its amplitude. While VEP differences in ocular dominance experiments are typically interpreted as changes in synaptic strength, this may not be entirely correct if NOR plasticity and latency are also changing.

Activity-dependent changes in NOR in other contexts suggest that just as synaptic plasticity operates through different mechanisms in association with development, pathology, physiological regulation, and experience-dependent modification, NOR plasticity also occurs in distinct contexts, operating through different cellular mechanisms. Large changes in nodal gap length can be produced during NOR remodeling in development, in demyelinating disorders (Fu et al., 2011), after non-physiological intense stimulation, or stressful conditions. For example, nodal gap length is greatly increased in auditory nerve following hyperstimulation after prolonged intense sound (Tagoe et al., 2014), and large changes in NOR gap length are also increased after prolonged sleep deprivation (Bellesi et al., 2018). These changes increase the probability of conduction failure and would seem to reflect pathological responses.

The size of the experience-dependent changes in NOR gap length in the present studies, while tenths of a micron, is substantial relative to the approximately one-micron size of nodes of Ranvier. Changes in NOR gap length of this magnitude would adjust conduction velocity within the normal range to influence spike time arrival and physiological function (Dutta et al., 2018), rather than promoting conduction failure. Nodal gap plasticity of this scale is produced biologically by reversibly severing the attachment of myelin sheaths to the axon via the cell adhesion molecule neurofascin-155 in the paranodal region (Fields and Dutta, 2019). NOR gap changes in this size range have previously been reported following spatial learning (Cullen, 2021) and motor learning (Bacmeister et al., 2022). Nodes of Ranvier much longer than one micron can lead to severe impairment of impulse conduction as seen in demyelinating diseases (Kokubun et al., 2010).

## Conclusion

These new findings reveal a previously unknown contributor to experience-dependent plasticity operating through changes in action potential conduction velocity and myelin plasticity. The axon-specific plasticity is consistent with OMP theory. Other mechanisms that affect conduction velocity could cooperate in shortening the latency of spike time arrival in visual cortex from the deprived eye; however, changes in NOR morphology and spike time arrival in visual cortex after MD open a new avenue for investigation in ocular dominance plasticity. These findings advance our understanding of ocular dominance plasticity and, more broadly, in other neural circuits where synaptic modification is driven by the spike time arrival of axonal inputs converging onto a common postsynaptic target.

Additionally, these findings carry translational relevance for therapeutic strategies involving monocular visual disruption and disease. Occlusion therapy for amblyopia by patching the stronger eye decreases pVEP amplitude relative to the lazy eye. It has also been reported that the latency to pVEP decreases after prolonged eye patching, but the mechanism is unclear (Lunghi et al., 2015). The shorter NOR gaps and shorter latency of pVEP from the deprived eye found in the present study could contribute to this effect. The findings highlight the importance of temporal synchrony from the two eyes to binocular neurons in visual disorders and their treatment. Temporal synchronization between the two eyes is known to be impaired in amblyopia. A recent study reports that providing a time-shifted stimulus to the eyes, re-synchronizes the ocular input and improves visual function (Eisen-Enosh et al., 2023). The possibility that conduction velocity in retinal ganglion cell axons could change by activity-dependent plasticity provides a new aspect to the field.

## Acknowledgements

We thank Louis Dye in the NICHD Microscopy Imaging Center for electron microscopy service, and Gregg Whitlock for contribution to graphical art in figures.

## Author contributions

Conceptualization: RDF

Investigation: RDF, MKF, WCF, ENS, PRL, MJ

Project administration: RDF, WL, JSB

Funding acquisition: RDF, WL, JSB Writing: RDF, MKF, WCH, ENS, PRL

Review and editing: RDF, MKF, WCH, ENS, PRL, MJ, WL, JSB

## Data and materials availability

Available upon request.

